# A *KARRIKIN INSENSITIVE2* paralog in lettuce mediates highly sensitive germination responses to karrikinolide

**DOI:** 10.1101/2021.10.13.464162

**Authors:** Stephanie E. Martinez, Caitlin E. Conn, Angelica M. Guercio, Claudia Sepulveda, Christopher J. Fiscus, Daniel Koenig, Nitzan Shabek, David C. Nelson

## Abstract

Karrikins (KARs) are chemicals in smoke that can enhance germination of many plants. *Lactuca sativa cv*. Grand Rapids (lettuce), germinates in the presence of nanomolar karrikinolide (KAR_1_). We found that lettuce is much less responsive to KAR_2_ or a mixture of synthetic strigolactone analogs, *rac*-GR24. We investigated the molecular basis of selective and sensitive KAR_1_ perception in lettuce. The lettuce genome contains two copies of *KARRIKIN INSENSITIVE2* (*KAI2*), a receptor that is required for KAR responses in *Arabidopsis thaliana. LsKAI2b* is more highly expressed than *LsKAI2a* in dry achenes and during early stages of seed imbibition. Through cross-species complementation assays in Arabidopsis we found that *LsKAI2b* confers robust responses to KAR_1_, but *LsKAI2a* does not. Therefore, LsKAI2b likely mediates KAR_1_ responses in lettuce. We compared homology models of the ligand-binding pockets of KAI2 proteins from lettuce and a fire follower, *Emmenanthe penduliflora*. This identified pocket residues 96, 124, 139, and 161 as candidates that influence the ligand-specificity of KAI2. Further support for the significance of these residues was found through a broader comparison of pocket residue conservation among 324 asterid KAI2 proteins. We tested the effects of substitutions at these four positions in *Arabidopsis thaliana KAI2* and found that a broad array of responses to KAR_1_, KAR_2_, and *rac*-GR24 could be achieved.

## INTRODUCTION

Plants use several strategies for regeneration in the post-fire environment, including regrowth from surviving tissue (e.g. epicormic buds), physical release of seeds (e.g. serotiny), and germination from the soil seed bank. The ecological significance of post-fire germination is perhaps best illustrated by fire ephemerals (or, pyroendemics), short-lived plants that in some cases emerge only in the first one or two years after a fire. However, more than 1200 plant species broadly distributed throughout the angiosperms show positive germination responses to aerosol smoke or smoke-water solutions. Searches for germination regulators among the thousands of compounds present in smoke have yielded a number of stimulants such as karrikins, glyceronitrile, and NO_2_, as well as inhibitors such as trimethylbutenolide and related furanones (Keeley and Fotheringham, 1997; Flematti et al., 2004; Light et al., 2010; Flematti et al., 2011; Nelson et al., 2012; Burger et al., 2018; Keeley and Pausas, 2018).

Karrikins (KARs) are a class of butenolide molecules found in smoke and biochar that are produced by pyrolysis of carbohydrates (Flematti et al., 2004; Kochanek et al., 2016). KARs were discovered through bioassay-guided fractionation of smoke-water. This approach used several species that show sensitive responses to smoke-water, including *Lactuca sativa cv*. Grand Rapids (Asterales; common name lettuce) and the Australian fire-followers *Conostylis aculeata* (Commelinales) and *Stylidium affine* (Asterales), as biological readouts for the presence of germination stimulants. Karrikinolide (KAR_1_), the first karrikin to be identified, enhanced germination of these species at concentrations below 1 nM. (Flematti et al., 2004; van Staden et al., 2004) At least six KARs are found in smoke (Flematti et al., 2009). KAR_1_ is presumed to be the most potent karrikin for many plants (Flematti et al., 2007; Sun et al., 2020). However, *Arabidopsis thaliana* is more sensitive to KAR_2_ than KAR_1_ (Nelson et al., 2009; Nelson et al., 2010)

KAR responses in plants are mediated by an α/β-hydrolase protein, KARRIKIN INSENSITIVE2 (KAI2)/HYPOSENSITIVE TO LIGHT (HTL) (Waters et al., 2012). Upon activation, KAI2/HTL associates with the F-box protein MORE AXILLARY GROWTH2 (MAX2)/DWARF3 (D3) and a subset of proteins in the SUPPRESSOR OF MAX2 1 (SMAX1)-LIKE (SMXL) family, SMAX1 and SMXL2. MAX2 acts within an SCF-type E3 ubiquitin ligase complex to target SMAX1 and SMXL2 for proteasomal degradation (Stanga et al., 2013; Stanga et al., 2016; Khosla et al., 2020; Zheng et al., 2020; Wang et al., 2020b). SMAX1, and presumably SMXL2, associate with the transcriptional co-repressors TOPLESS (TPL) and TPL-related (TPR) through an EAR motif (Soundappan et al., 2015). Therefore, degradation of SMAX1 and SMXL2 is thought to relieve transcriptional repression and initiate downstream growth responses. In addition to promoting seed germination, the KAR signaling pathway enhances seedling photomorphogenesis, root hair density, root hair elongation, and stress tolerance; suppresses mesocotyl elongation in the dark; and enables symbiotic interactions with arbuscular mycorrhizal fungi (Nelson et al., 2010; Waters et al., 2012; Gutjahr et al., 2015; Li et al., 2017; Wang et al., 2018; Villaécija-Aguilar et al., 2019; Carbonnel et al., 2020a; Choi et al., 2020; Li et al., 2020; Shah et al., 2020; Zheng et al., 2020; Carbonnel et al., 2020b).

KAI2 is an ancient paralog of the strigolactone (SL) receptor DWARF14 (D14)/DECREASED APICAL DOMINANCE2 (DAD2)/RAMOSUS3 (RMS3) (Hamiaux et al., 2012; de Saint Germain et al., 2016; Yao et al., 2016). SLs are butenolide molecules like KARs, but have altogether different molecular structures, sources, and functions in plants. Canonical SLs consist of a tricyclic ABC-ring connected by an enol-ether bond to a butenolide D-ring in the 2’*R* stereochemical configuration. Noncanonical SLs are similar, but lack cyclized ABC rings (Al-Babili and Bouwmeester, 2015; Yoneyama et al., 2018). SLs are carotenoid-derived plant hormones that regulate axillary bud outgrowth (i.e. shoot branching or tillering), leaf senescence, cambium development, and drought tolerance, among other developmental processes (Gomez-Roldan et al., 2008; Umehara et al., 2008; Agusti et al., 2011; Ueda and Kusaba, 2015; Li et al., 2020). SLs are also exuded into the soil, particularly under low N or P conditions, which promotes beneficial symbiotic interactions with arbuscular mycorrhizal fungi (Akiyama et al., 2005; Al-Babili and Bouwmeester, 2015; Yoneyama et al., 2018). Obligate parasitic plants in the Orobanchaceae, such as witchweeds (*Striga* spp.) and broomrapes (*Orobanche, Phelipanche* spp.) have evolved the ability to use very low levels of SLs in the rhizosphere as germination cues that indicate the presence of a potential host (Bouwmeester et al., 2021; Nelson, 2021). SL signaling is highly similar to KAR signaling. Upon activation by SL, D14 works with SCF^MAX2^ to target a different subset of SMXL proteins (SMXL6, SMXL7, and SMXL8 in Arabidopsis, or the orthologous DWARF53 protein (D53) in rice) for proteasomal degradation (Jiang et al., 2013; Zhou et al., 2013; Soundappan et al., 2015; Wang et al., 2015; Liang et al., 2016; Yao et al., 2016). Like SMAX1 and SMXL2, DWARF53 and its orthologs have one or more conserved EAR motifs and interact with TPL/TPR proteins (Jiang et al., 2013; Soundappan et al., 2015; Wang et al., 2015). D53 regulates downstream gene expression in monocots through interaction with SQUAMOSA-PROMOTER BINDING PROTEIN-LIKE (SPL) transcription factors (Liu et al., 2017; Song et al., 2017). SMXL6 is also likely to work with transcription factors, but can bind DNA to regulate transcription directly as well (Wang et al., 2020a).

KAI2 proteins in plants collectively perceive a diverse range of signals. In Arabidopsis, KAI2 mediates responses to GR24^*ent*-5DS^, a synthetic SL analog that has a D-ring in an unnatural 2’*S* configuration, as well as KARs (Waters et al., 2012; Scaffidi et al., 2014). There is substantial biochemical evidence that KAI2 can bind KAR_1_, which in combination with genetic evidence has led to the reasonable conclusion that KAI2 is a KAR receptor (Guo et al., 2013; Kagiyama et al., 2013; Toh et al., 2014; Xu et al., 2016; Lee et al., 2018; Xu et al., 2018; Bürger et al., 2019). However, it now seems more likely that KAI2 perceives an unknown KAR metabolite(s) (Waters et al., 2015; Xu et al., 2018). Differential scanning fluorimetry (DSF) assays show that KAI2 undergoes thermal destabilization *in vitro* in the presence of GR24^*ent*-5DS^, but not KAR_1_ or KAR_2_ (Waters et al., 2015). Similarly, *rac*-GR24 or GR24^*ent*-5DS^, but not KAR_1_, promotes protein-protein interactions between KAI2 and MAX2, SMAX1, or SMXL2 (Xu et al., 2018; Khosla et al., 2020; Wang et al., 2020b). Therefore, unmodified KARs probably cannot activate KAI2 directly, but GR24^*ent*-5DS^ can. Putatively, KAI2 also perceives an undiscovered, endogenous signal in plants known as KAI2 ligand (KL) (Conn and Nelson, 2015). Evidence for KL include *kai2* and *max2* mutant phenotypes that are opposite to the effects of KAR treatment which are not observed in SL biosynthesis or signaling mutants (Nelson et al., 2011; Waters et al., 2012). In addition, highly <u>c</u>onserved *KAI2* paralogs (KAI2c) in root parasitic plants can rescue an Arabidopsis *kai2* mutant but do not respond to KARs, suggesting they sense another signal in plants (Conn et al., 2015; Conn and Nelson, 2015). KAI2 has undergone an atypical degree of gene duplication in the Orobanchaceae (Lamiales), resulting in a parasite-specific clade of fast-evolving, <u>d</u>ivergent KAI2 paralogs (KAI2d) that can perceive different SLs and, in one case, isothiocyanates (Conn et al., 2015; Toh et al., 2015; Tsuchiya et al., 2015; de Saint Germain et al., 2021). D14 itself is another example of a SL receptor that was derived from KAI2 (Bythell-Douglas et al., 2017). Finally, a KAI2 representative from a grade that has undergone an intermediate level of purifying selection, *Striga hermonthica* KAI2i (*ShKAI2i*), confers KAR-specific responses and only weakly rescues Arabidopsis *kai2* (Conn et al., 2015; Conn and Nelson, 2015).

These observations raise the question of how different ligand specificities have evolved in KAI2 proteins, enabling some plants to gain beneficial traits such as post-fire germination or host-induced germination. We set out to determine the basis of highly sensitive germination responses to KAR in lettuce, which we reasoned might give clues to how some species have adapted to fire-prone ecosystems.

## RESULTS

### Lettuce achenes are more sensitive to KAR_1_ than KAR_2_

We first tested whether lettuce achenes have different sensitivity to KAR_1_, KAR_2_, and *rac*-GR24, a racemic mixture of GR24^*ent*-5DS^ and its 2’*R*-configured enantiomer GR24^5DS^ (Figure 1A). Prior work had suggested that KAR_1_ is a more effective stimulant of lettuce germination than KAR_2_, but the compounds were tested in separate experiments, limiting a direct comparison (Flematti et al., 2007). We found that 1 µM KAR_1_ and KAR_2_ treatments induced ∼100% germination of lettuce in the dark, compared to ∼40% germination for mock-treated achenes (Figure 1B). 1 µM *rac*-GR24 also stimulated lettuce germination, but was less effective than either KAR. To determine whether KAR_1_ or KAR_2_ is more potent, we evaluated the effects of a range of KAR_1_ and KAR_2_ concentrations on lettuce germination. KAR_1_ induced nearly complete germination at 1 nM and higher concentrations (Figure 1C). By contrast, 1 nM and 10 nM KAR_2_ did not enhance germination, and 100 nM KAR_2_ had an intermediate effect compared to 1 µM KAR_2_.

**Figure 1.**
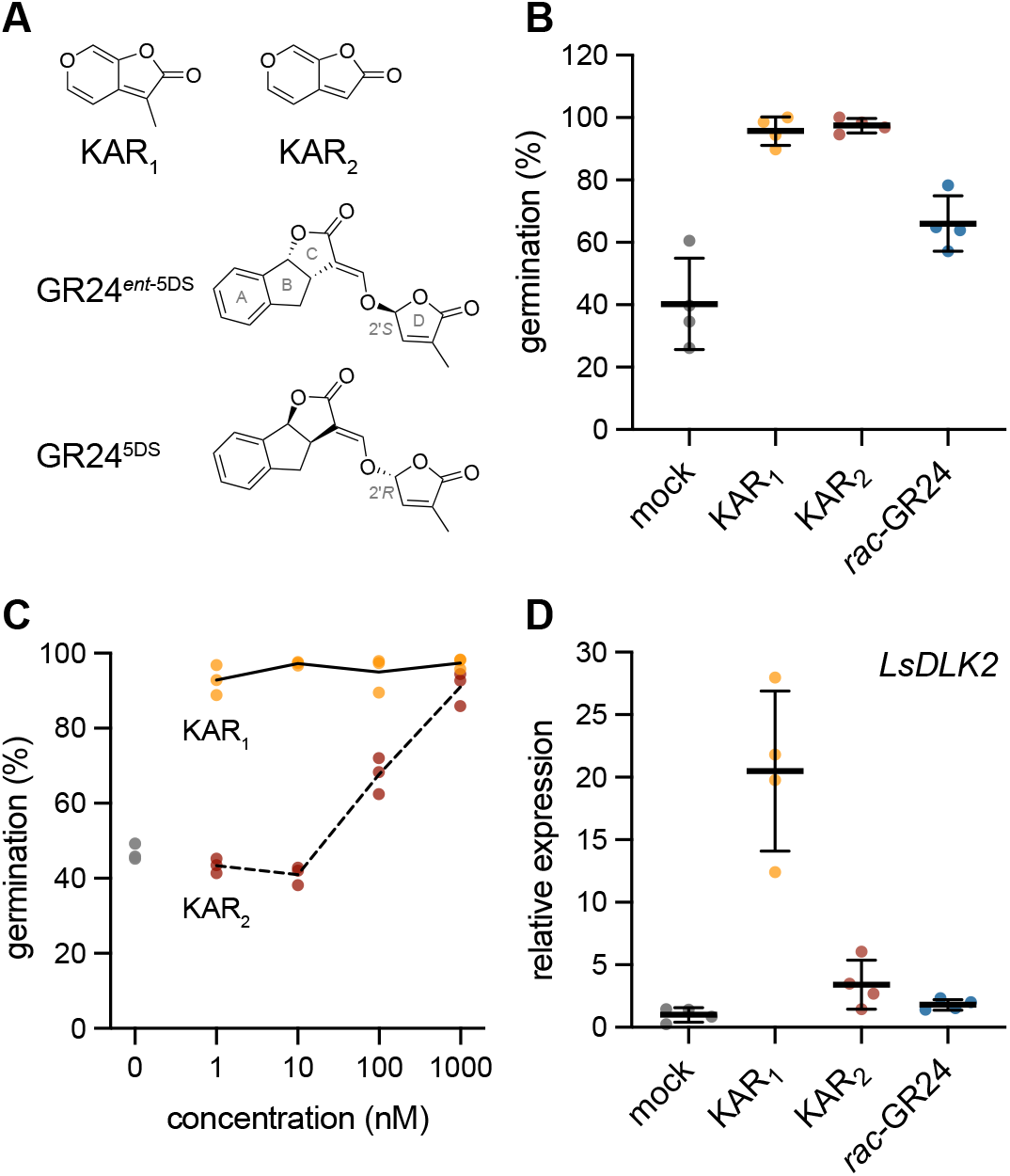
Lettuce achenes are highly sensitive to KAR_1_. Structures of KAR_1_, KAR_2_, GR24^*ent*-5DS^, and GR24^5DS^. GR24^5DS^ is a strigolactone analog that mimics the stereochemical configuration of the SL 5-deoxystrigol. GR24^*ent*-5DS^ is an enantiomer of GR24^5DS^ that has a methylbutenolide D-ring in 2’*S* configuration, which is not found in natural SLs. Lettuce germination in the presence of 0.1% (v/v) acetone or 1 µM KAR_1_, KAR_2_, or *rac*-GR24. Achenes were incubated 1 h in darkness, followed by a pulse of far-red light for 10 minutes, and the remaining 48 h in darkness at 20°C. n=4 replicates of approximately 50-70 achenes each; mean ± SD. C) Lettuce germination in the presence of a range of KAR_1_ and KAR_2_ concentrations after 1 h in darkness, followed by a pulse of far-red light for 10 minutes, and the remaining 48 h in darkness at 20°C. n=3 replicates of approximately 50-60 achenes each. D) qRT-PCR analysis of *LsDLK2* expresion relative to *LsACT* (actin) in lettuce achenes imbibed with 0.1% (v/v) acetone or 1 µM KAR_1_, KAR_2_, or *rac*-GR24 for 24 h in darkness at 20°C. n=4 pools of achenes; mean ± SD. Values re-scaled to relative *LsDLK2* expression in mock-treated achenes.

As an additional test of lettuce achene responses to KAR_1_, KAR_2_, and *rac*-GR24, we examined how each treatment affected the expression of *D14-LIKE2* (*DLK2*). *DLK2* is an ancient paralog of *KAI2* and *D14* that serves as a transcriptional marker of KAR/KL signaling in diverse angiosperms such as *Arabidopsis thaliana, Brassica tournefortii, Oryza sativa* (rice), and *Lotus japonicus* (Waters et al., 2012; Sun et al., 2016; Sun et al., 2020; Carbonnel et al., 2020b). In achenes imbibed 24 h in the dark with KAR or *rac*-GR24 treatments, *LsDLK2* (Lsat_1_v5_gn_8_94781) transcripts were induced approximately 20-fold by 1 µM KAR_1_ compared to mock treatment (Figure 1D). *LsDLK2* transcripts were increased ∼4-fold by 1 µM KAR_2_ and ∼2-fold by 1 µM *rac*-GR24 compared to mock treatment, but these changes were not statistically significant (p > 0.05, Dunnett’s T3 multiple comparisons test). Altogether, these data indicate that lettuce achenes are much more sensitive to KAR_1_ than KAR_2_ or *rac*-GR24.

### Lettuce has two *KAI2* paralogs that have differential expression in seed

We hypothesized that altered ligand-specificity or -affinity of a KAI2 protein(s) may underlie the high sensitivity to KAR_1_ observed in lettuce. A BLAST search of the lettuce draft genome revealed two putative *KAI2* orthologs that we designated *LsKAI2a* and *LsKAI2b* (Table S1). We performed a phylogenetic analysis to determine whether either of these genes are *KAI2i*-type, which had previously been associated with selective responses to KAR_1_ (Conn et al., 2015; Conn and Nelson, 2015). We found that neither *LsKAI2* paralog grouped with the Lamiid *KAI2i* grade (Figure 2). Instead, LsKAI2a and LsKAI2b form a monophyletic sister group to the Lamiid *KAI2c* clade, suggesting that they emerged from a gene duplication that occurred after the divergence of the Lamiids and the Campanulids.

**Figure 2.**
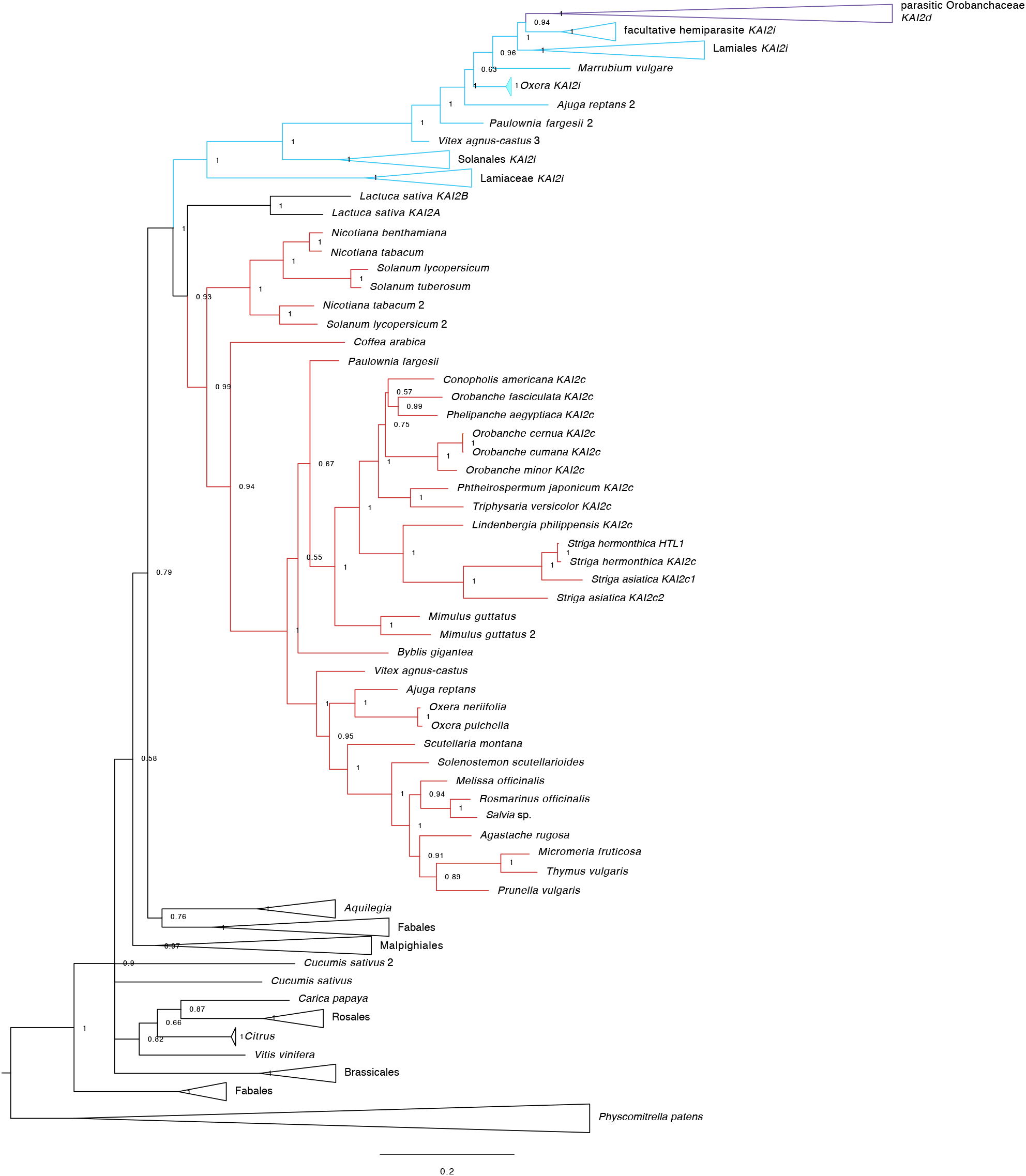
Lettuce *KAI2* genes group with the conserved *KAI2c* clade. Bayesian phylogeny of *KAI2* genes in dicots. Sequences from lamiids fall into the conserved (*KAI2c*, red), intermediate (*KAI2i*, blue), and divergent (*KAI2d*, purple; parasite-specific) clades that were previously described (Conn et al., 2015).

We examined whether *LsKAI2a* and *LsKAI2b* are expressed at different levels in achenes, as it might highlight one gene as a more likely candidate for mediating KAR_1_ responses during germination. In dry, unimbibed achenes, *LsKAI2b* transcripts were ∼5-fold more abundant than *LsKAI2a* transcripts. *LsKAI2b* transcript abundance progressively declined after 6 h and 24 h of imbibition, whereas *LsKAI2a* rose slightly after 24 h imbibition (Figure 3). This suggested that LsKAI2b protein may be more abundant than LsKAI2a during the early stages of imbibition.

**Figure 3.**
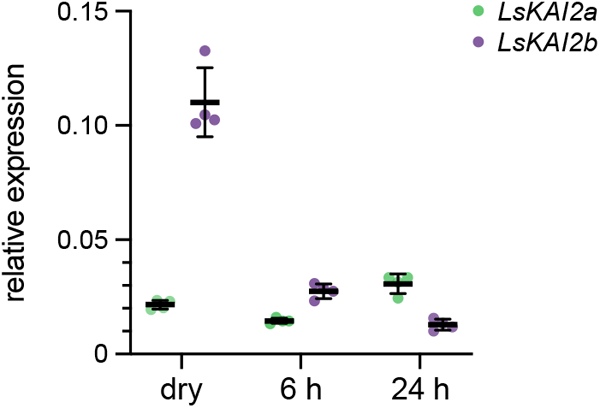
*LsKAI2b* transcripts are more abundant than *LsKAI2a* during early imbibition. qRT-PCR analysis of *LsKAI2a* and *LsKAI2b* expression relative to *LsACT* in lettuce achenes that were un-imbibed (dry), or imbibed in water for 1 h in darkness, followed by 10 min in far-red light, and the remaining time to 6 or 24 h in darkness at 20°C. n=4 pools of achenes; mean ± SD.

### *LsKAI2b* confers KAR_1_ responses to Arabidopsis seedlings

We then set out to determine whether LsKAI2a and LsKAI2b proteins have different ligand preferences. Because KAR metabolites and the endogenous KL remain unknown, it was not possible to test receptor-ligand affinities *in vitro*. Instead, we used cross-species complementation assays to investigate the responses of LsKAI2a and LsKAI2b to KARs and *rac*-GR24. We cloned *LsKAI2a* and *LsKAI2b* genes (coding sequence including intron) into a plant transformation vector that drives transgene expression from an *Arabidopsis thaliana KAI2* (AtKAI2) promoter. We generated homozygous transgenic lines for *AtKAI2p:LsKAI2A* and *AtKAI2p:LsKAI2B* in an Arabidopsis *d14 kai2* background, which does not respond to KARs or *rac*-GR24. As a control, we tested *d14 kai2* lines transgenic for an Arabidopsis *KAI2* coding sequence.

We tested the inhibitory effects of 1 µM KAR_1_, KAR_2_, and *rac*-GR24 on hypocotyl elongation of seedlings grown under continuous red light (Figure 4). This assay provides a useful alternative to Arabidopsis germination tests, which are often challenging to perform consistently due to variable and labile seed dormancy. As expected, *AtKAI2p:AtKAI2* rescued the elongated hypocotyl phenotype of *d14 kai2* and restored responses to KAR_1_, KAR_2_, and *rac*-GR24. KAR_2_ caused a stronger reduction in hypocotyl elongation than KAR_1_, as observed in wild-type (wt) Col-0 Arabidopsis seedlings. Responses to *rac*-GR24 were partially reduced compared to wt due to the lack of *D14*-mediated signaling. The *LsKAI2a* transgene had mixed effects in different transgenic lines. All lines had reduced hypocotyl length under mock-treated conditions compared to *d14 kai2*, suggesting rescue of KL response. Only the line with the strongest *LsKAI2a* expression showed a response to KAR_1_ or KAR_2_, and this was weak compared to KAR responses in *AtKAI2* transgenic lines (Figure 4, S1A). Inhibition of hypocotyl elongation by *rac*-GR24 was also weak, and was only observed in two transgenic lines. By contrast, *LsKAI2b* conferred a strong and specific response to KAR_1_. The degree of hypocotyl growth inhibition by KAR_1_ in *LsKAI2b* lines exceeded that observed in wt and *AtKAI2* transgenic lines, despite lower levels of *LsKAI2b* expression compared to *AtKAI2* (Figure 4, S1A). KAR_2_ did not affect hypocotyl elongation of *AtKAI2p:LsKAI2b* lines, and *rac*-GR24 had inconsistent and comparatively weak effects. Hypocotyl elongation under mock-treated conditions was only reduced in one transgenic line, which had the highest *LsKAI2b* expression. In terms of rescue of the reduced expression of *DLK2* in *d14 kai2* seedlings, neither *LsKAI2a* or *LsKAI2b* were as effective as *AtKAI2* (Figure S1B).

**Figure 4.**
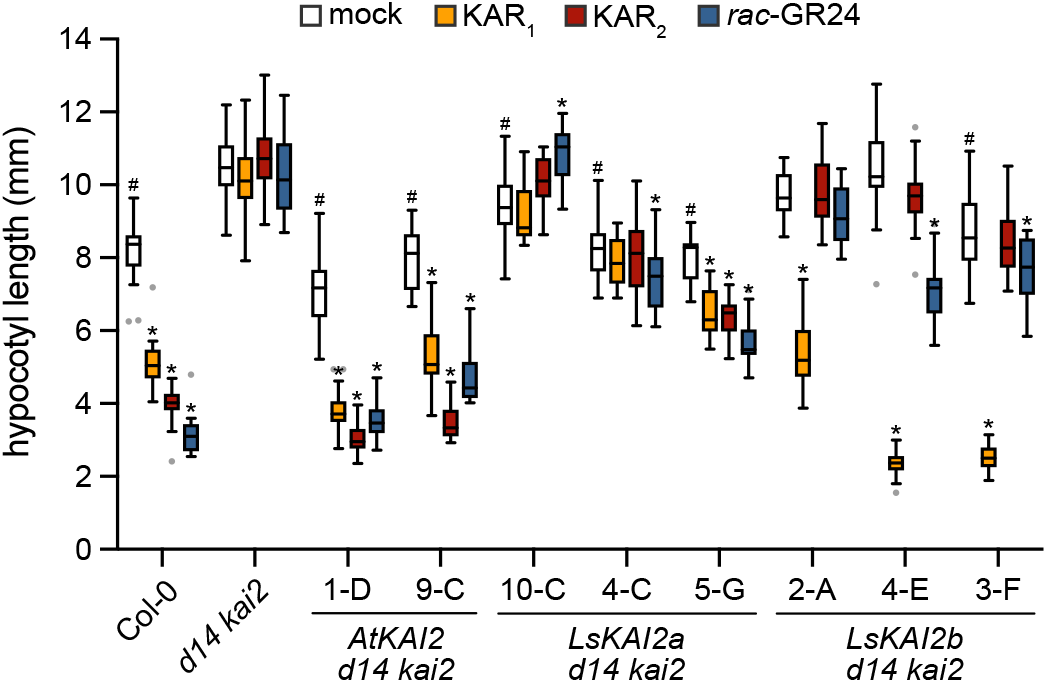
An LsKAI2b transgene confers strong KAR_1_ responses to Arabidopsis seedlings. Hypocotyl length of 5-d-old *Arabidopsis thaliana* seedlings grown under red light on 0.5x MS media supplemented with 0.1% (v/v) acetone or 1 µM KAR_1_, KAR_2_, or *rac*-GR24. n=20 seedlings. Box plots indicate median and quartiles with Tukey’s whiskers. Gray dots indicate outlier data beyond Tukey’s whiskers. *, p<0.01, Dunnett’s multiple comparisons test, treatment versus mock comparison within each line. ^#^, p<0.01, Dunnett’s multiple comparisons test, comparison to *d14 kai2*, mock-treated samples only.

We compared the responses of wt and *AtKAI2p:LsKAI2b d14 kai2* seedlings to a range of KAR_1_ concentrations (Figure S1C). We found that 100 nM and higher concentrations of KAR_1_ caused a reduction in hypocotyl elongation of wt seedlings. In an *LsKAI2b* transgenic line, however, 1 nM KAR_1_ was sufficient to cause a similar response (Figure S1C). In the presence of 1 µM KAR_1_, hypocotyl elongation was inhibited 78% in the *LsKAI2b* transgenic line compared to 38% inhibition in wt. As the expression of *LsKAI2b* was at least two-fold lower than endogenous *KAI2* in wt Arabidopsis, this suggests that LsKAI2b is highly effective at transducing KAR_1_ responses (Figure S1A).

### Structural differences in the LsKAI2b pocket may influence KAR_1_ sensitivity

Although LsKAI2a and LsKAI2b proteins share 84% amino acid identity and 94% similarity, LsKAI2b is uniquely able to confer highly sensitive KAR_1_ responses to *Arabidopsis thaliana*. We investigated which amino acid differences might alter the ligand specificity and/or affinity of LsKAI2b. To predict the overall structures and ligand-binding pocket morphologies of LsKAI2a and LsKAI2b, we generated protein structure homology models using Phyre2 with ShKAI2iB (PDB structure 5DNW) as a template (Kelley et al., 2015; Xu et al., 2016). Of the 43 amino acid differences between LsKAI2a and LsKAI2b, seven are in pocket-defining residues: V/M96, Y/F124, Y/F134, D/E138E, L/V139, M/I146, A/V161 (Figure 5A,B; for consistency, equivalent AtKAI2 position numbers are used here and below). The differences at these positions are predicted to substantially enlarge the volume of the pocket in LsKAI2b relative to LsKAI2a (Figure 5A,B). LsKAI2b has a pocket volume of 189 Å^3^, compared to 126 Å^3^ for LsKAI2a and 126 Å^3^ for AtKAI2 (PDB structure 5Z9G) (Lee et al., 2018). Particularly notable differences that affect pocket shape in lettuce KAI2 proteins are found among the residues that surround α-helix 4; these residues are bulkier and more of them are positively charged in LsKAI2a than in LsKAI2b (Figure 5A-B).

**Figure 5.**
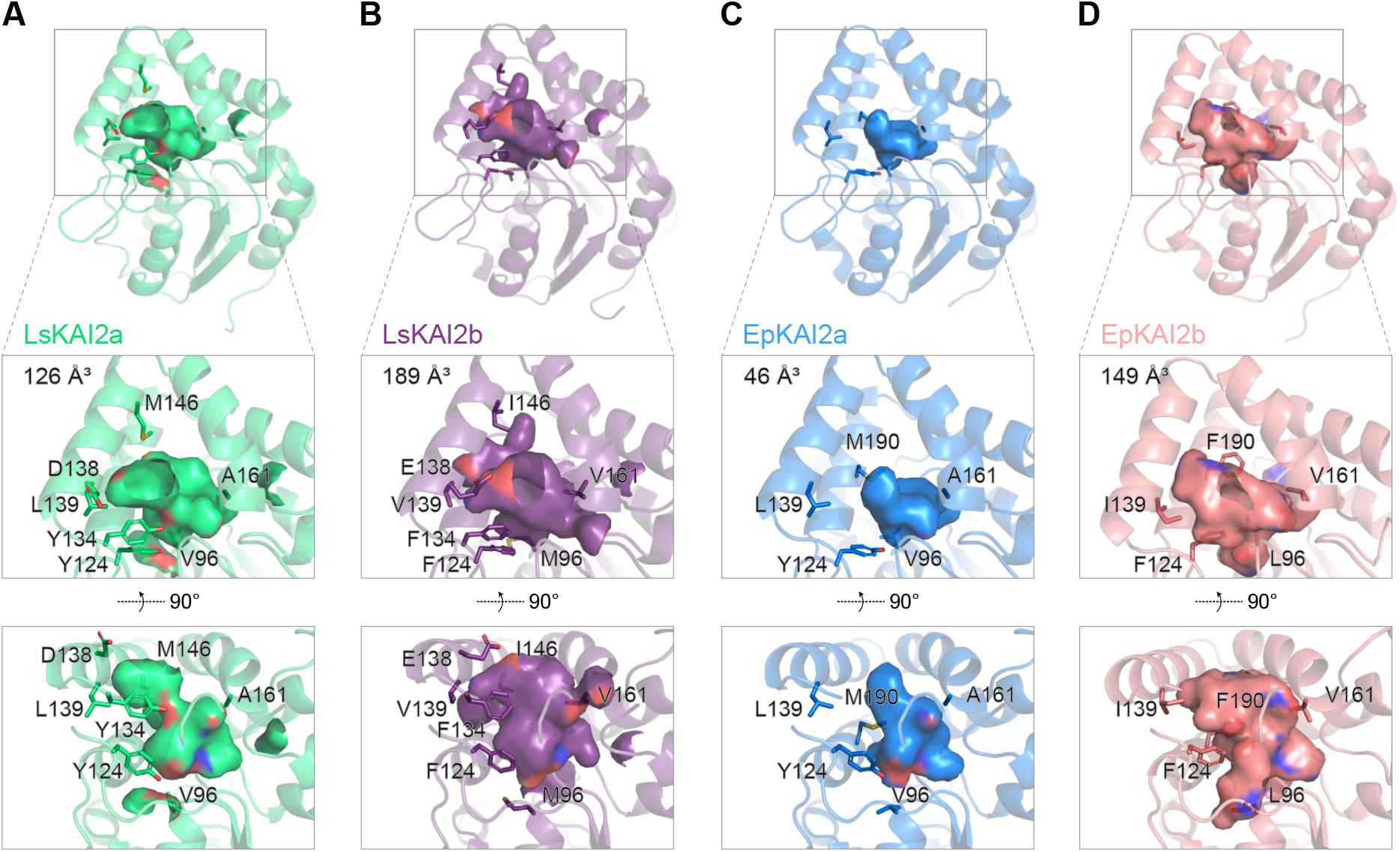
KAI2b proteins in lettuce and *E. penduliflora* have enlarged ligand-binding pockets. Homology models of A) LsKAI2a, B) LsKAI2b, C) EpKAI2a, and D) EpKAI2b. Hydrophobic cavities and their volumes are shown. Pocket residues that differ between KAI2a and KAI2b in each species are indicated.

We hypothesized that similar changes to the volume or chemical properties of the KAI2 ligand-binding pocket as those found in lettuce might have occurred in other species that have evolved sensitive responses to KAR_1_. The Californian chaparral species, *Emmenanthe penduliflora* (Boraginales; common name, “whispering bells”) is a smoke-responsive annual that primarily emerges in post-fire sites. Its germination can be triggered by nitric oxides (e.g. NO_2_) from smoke as well as by <10 nM KAR_1_ (Keeley and Fotheringham, 1997; Flematti et al., 2004; Flematti et al., 2007). To investigate *KAI2* evolution in this fire-follower, we generated a *de novo* transcriptome assembly from RNA extracted from seedlings. We identified two *KAI2* coding sequences (Table S1). The predicted EpKAI2a and EpKAI2b protein sequences are 74% identical. Among the 70 amino acid differences, five are in pocket-defining residues (V/L96, Y/F124, L/I139, A/V161, M/F190). As with lettuce KAI2 proteins, the differences at these five positions are predicted to substantially enlarge the pocket volume of EpKAI2b compared to EpKAI2a (Figure 5C,D). The root mean square deviation (RMSD) for LsKAI2a and EpKAI2a models is 0.093 Å (Figure S2). By contrast, comparisons of LsKAI2b and LsKAI2a, EpKAI2b and EpKAI2a, and LsKAI2b and EpKAI2b indicate larger RMSD values ranging from 0.42 to 0.63 Å (Figure S2). These models suggest that EpKAI2b is a more likely candidate for a KAR_1_-specific receptor than EpKAI2a, which seems likely to have ligand preferences that are similar to LsKAI2a. Notably, four of the pocket-defining positions that distinguish EpKAI2a and EpKAI2b overlapped with those that distinguish LsKAI2a and LsKAI2b; specifically, positions 96, 124, 139, and 161.

### Conserved pocket residue changes among two major groups of asterid KAI2 paralogs

We also compared the amino acid sequences of the parasitic plant proteins *Phelipanche aegyptiaca* KAI2c (PaKAI2c) and *Striga hermonthica* KAI2c (ShKAI2c) to ShKAI2i, which confers sensitive responses to KAR_1_ to Arabidopsis (Conn et al., 2015; Conn and Nelson, 2015; Toh et al., 2015). Among the eight total positions that distinguish KAI2a and KAI2b pockets in either lettuce or *E. penduliflora*, seven substitutions (V96L, Y124F, E138D, L139V, M146I, C161V, and A/L190F) were observed in ShKAI2i relative to PaKAI2c and ShKAI2c (Figure S3). This revealed several pocket residue substitutions in KAI2b/KAI2i proteins that were consistent in lettuce, *E. penduliflora*, and *Striga hermonthica*: V96L/M, Y124F, L139V/I, and A/C161V. If EpKAI2b is selectively responsive to KAR_1_ like LsKAI2b and ShKAI2i, we hypothesized that these shared changes may cause their KAR_1_ specificity.

To investigate whether similar differences occur at these positions among KAI2 paralogs in other asterids, we performed an in-depth examination of KAI2 protein sequences in *de novo* transcriptome assemblies that had been generated by the One Thousand Plants (1KP) consortium (One Thousand Plant Transcriptomes Initiative, 2019). Through reciprocal BLAST searches we identified 334 KAI2 protein sequences from 199 species (Table S2). As suggested by the Y124F substitution that was shared by LsKAI2b, EpKAI2b, and ShKAI2i, we found that asterid KAI2 proteins could be split into two major groups based upon Tyr or Phe identity at position 124. Only 12 of the 334 KAI2 proteins (3.6%) did not have Y124 or F124 residues.

Almost all species (188 of 199, 94.5%) had at least one Y124-type KAI2 paralog. By contrast, F124-type KAI2 paralogs were found in less than half of the species (88 of 199, 44.2%; Figure 6A; Table S3). For 94 species, only one KAI2 was identified; for 85 of these species (90.4%), this protein was Y124-type and for 6 species (6.4%) it was F124-type. Although some *KAI2* genes are likely to have been missing from the *de novo* transcriptome assemblies (e.g. due to inadequate sequencing depth or RNA sampling), the disparity in these distributions suggests that plants require Y124-type KAI2 proteins, while F124-type KAI2 proteins may have more auxiliary functions. Notably, an F124-type KAI2 was not observed in any of the 36 Asterales transcriptomes surveyed (Figure 6A). This suggested that the emergence of an F124-type KAI2 (i.e. LsKAI2b) in lettuce may have occurred independently within the Asterales lineage. By contrast, the presence of an F124-type KAI2 (i.e. EpKAI2b) in *E. penduliflora* is typical for the Boraginales.

**Figure 6.**
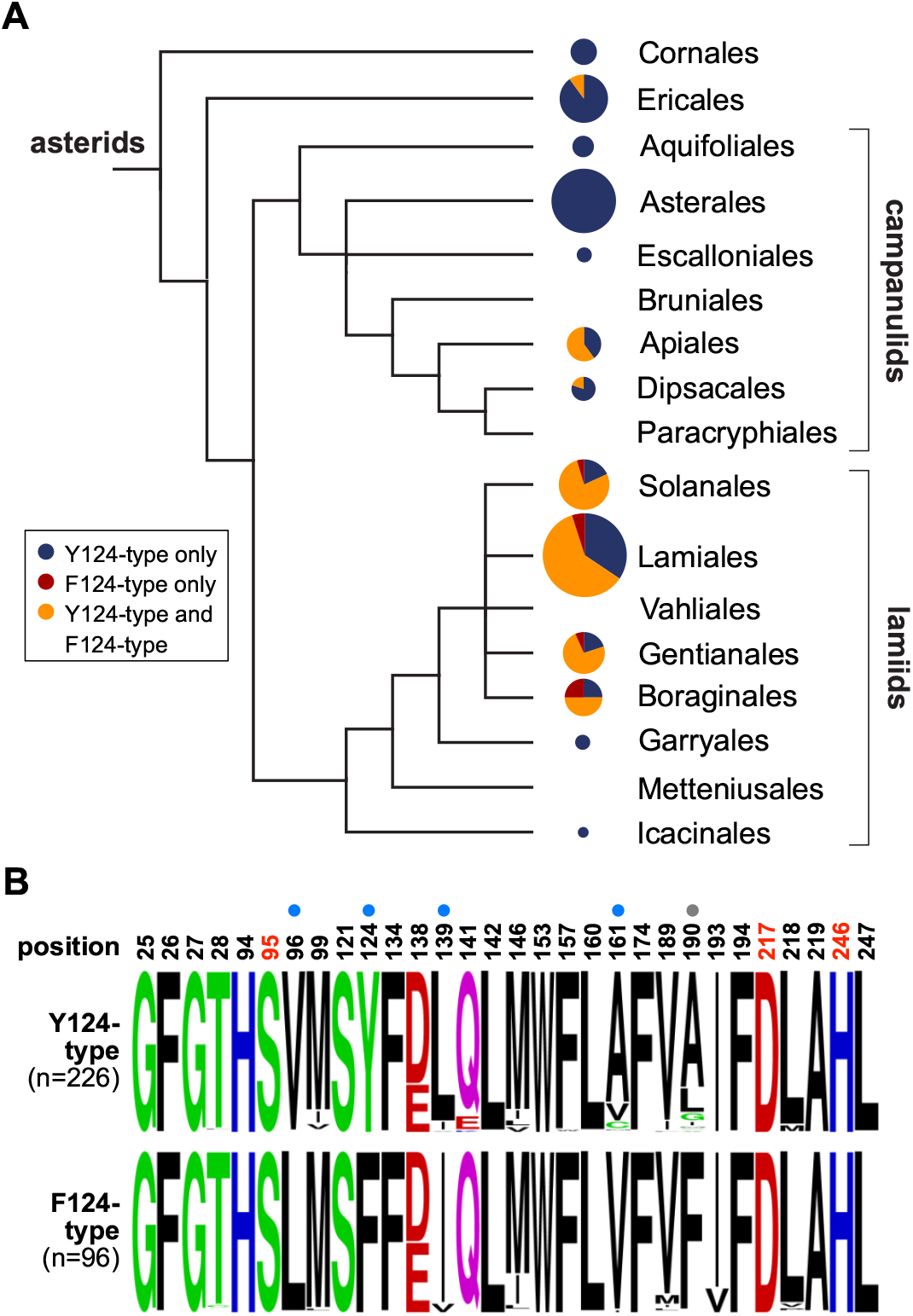
Two groups of asterid KAI2 have conserved differences at five pocket positions. A) Distribution of KAI2 types in asterids. Phylogeny adapted from Angiosperm Phylogeny Group (APG) IV system <u>(Chase et al., 2016)</u>. Pie charts indicate the proportion of species for which only Y124-type KAI2 (blue), only F124-type KAI2 (red), or both Y124-type and F124-type KAI2 (orange) were observed in *de novo* transcriptome assemblies from OneKP. The area of each pie chart is proportional to the number of species that were sampled from each order, from n=1 for Icacinales to n=60 for Lamiales. B) Frequency plots of amino acid composition in asterid KAI2 proteins at 30 positions that form the ligand-binding pocket. Asterid KAI2 proteins were split into two groups based upon Tyr or Phe amino acid identity at position 124. Dots above residues indicate candidates for ligand specificity-determining residues based upon amino acid conservation within and across the two groups. Blue dots indicate prioritized candidate positions. Position 190 was de-prioritized because LsKAI2b does not have a Phe190 residue but is sensitive to KAR_1_ nonetheless.

We identified 30 positions that define the KAI2 ligand-binding pocket. We examined amino acid conservation at these positions within the two major KAI2 groups in asterids. A high degree of conservation was observed within and across the two groups at 25 positions (Figure 6B). However, four positions in addition to 124 were well-conserved within each group and different between the groups. Positions 96, 124, 139, 161, and 190 stood out as candidates that might define ligand-specificity differences between the KAI2 groups. This broader analysis gave us reason to exclude positions 134, 138, and 146, which had been identified in LsKAI2a and LsKAI2b comparisons, from further consideration, as similar amino acid compositions were found among the two KAI2 groups. Position 190 was also de-prioritized, as LsKAI2b does not share the F190 identity of other F124-type KAI2, but nonetheless confers sensitive KAR_1_ responses (Figure S3).

### The impact of four pocket residues on ligand-specificity of KAI2

To investigate whether positions 96, 124, 139, and 161 influence ligand-specificity, we generated a series of substitutions in AtKAI2 proteins. AtKAI2 shares V96, Y124, L139, and A161 identities with LsKAI2a, EpKAI2a, and most asterid Y124-type KAI2 proteins. We mimicked asterid F124-type KAI2 proteins at these positions by creating quadruple and triple mutant combinations of V96L, Y124F, L139I, and A161V substitutions in AtKAI2. (The variants are annotated here in superscripts by amino acid identities at positions 96, 124, 139, and 161, respectively, with non-AtKAI2 substitutions underlined.) The *AtKAI2* variants were introduced into the Arabidopsis *d14 kai2* background and homozygous transgenic lines were tested for responses to KAR_1_, KAR_2_, and *rac*-GR24. We observed a range of ligand specificities among the variants. AtKAI2^V<u>FIV</u>^ was KAR_1_-specific, AtKAI2<u>^LYIV^</u> was KAR_1_- and KAR_2_-specific, AtKAI2<u>^LFIA^</u> showed reduced responses to KAR_1_ and KAR_2_, and the quadruple mutant AtKAI2<u>^LFIV^</u> was KAR_1_- and *rac*-GR24-specific (Figure 7A, S4). Interestingly, AtKAI2<u>^LFIV^</u> conferred a stronger response to KAR_1_ than triple-substituted variants or wt *AtKAI2*. AtKAI2<u>^LFLV^</u> may be KL-specific, as it did not confer consistent responses to any of the treatments but did rescue hypocotyl elongation under mock-treated conditions.

**Figure 7.**
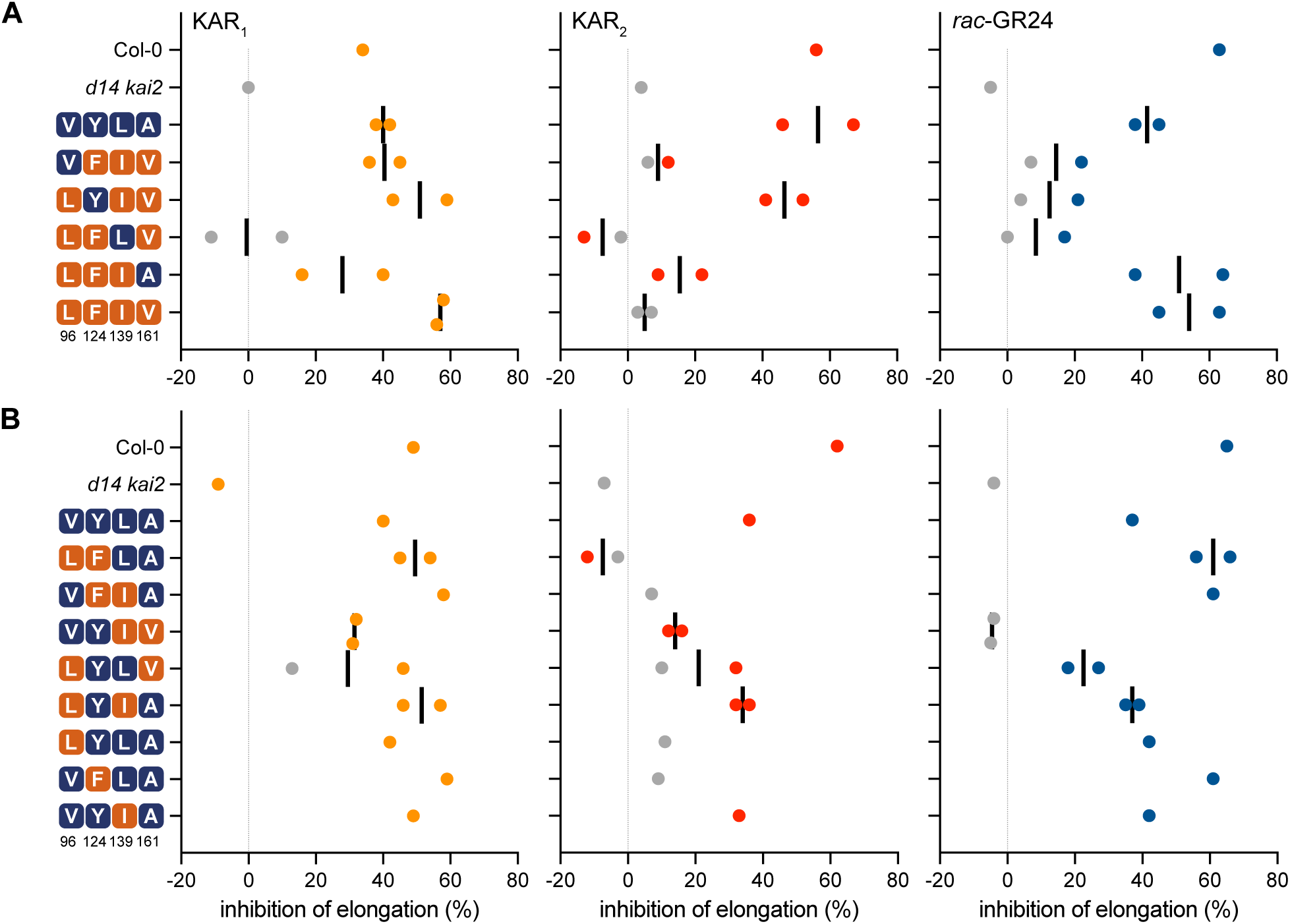
Pocket residues at positions 96, 124, 139, and 161 affect AtKAI2 ligand-specificity. Inhibition of hypocotyl elongation by KAR_1_, KAR_2_, or *rac*-GR24 in 5-d-old seedlings grown in red light for transgenic lines of *AtKAI2* variants with A) quadruple- and triple-, or B) double- and single-substitutions. All transgenic lines are in the *d14 kai2* double mutant background. Data are derived from hypocotyl length measurements shown in Figure S4. Each data point represents growth inhibition for a unique genetic line. Gray points indicate data that were not significantly different from mock-treated controls for each transgenic line.

We attempted to establish a relationship between specific amino acid changes in KAI2 and ligand specificity. We noted that KAR_2_ response was highly reduced or absent in AtKAI2^V<u>FIV</u>^, AtKAI2<u>^LFIA^</u>, and AtKAI2<u>^LFIV^</u> lines (Figure 7A, S4A). This suggested that Y124F, L139I, or a combination of both substitutions abolishes KAR_2_ perception. Responses to *rac*-GR24 were highly reduced in AtKAI2^V<u>FIV</u>^ and AtKAI2^LY<u>IV</u>^, which share L139I and A161V substitutions. This suggested that these positions may be relevant to *rac*-GR24 perception. However, it must be noted that AtKAI2<u>^LFIV^</u> also has L139I and A161V substitutions but remained responsive to *rac*-GR24 (Figure 7A, S4A). The shortest hypocotyls under mock-treated conditions, which may indicate KL responsiveness, were observed in AtKAI2<u>^LFIV^</u> transgenic lines (Figure S4A).

To investigate these hypotheses further, we examined the effects of three of the four single substitutions and five of the six possible double substitution combinations at positions 96, 124, 139, and 161 (Figure 7B, S4B). Among the double mutants, AtKAI2<u>^LFLA^</u> and AtKAI2^V<u>F</u>IA^ had similar effects; KAR_2_ response was lost, while KAR_1_ and *rac*-GR24 response remained. AtKAI2^VY<u>IV</u>^ lost responses to *rac*-GR24 and had reduced responses to KAR_2_ and, to a lesser extent, KAR_1_. AtKAI2^LYL<u>V</u>^ showed similar responses to KARs as AtKAI2^VY<u>IV</u>^, but was not as strongly affected in its *rac*-GR24 response. AtKAI2^LYIA^ conferred similar responses to wt AtKAI2. Among the single mutants, AtKAI2^LYLA^ had highly reduced responses to KAR_2_ and wt responses to KAR_1_ and *rac*-GR24. AtKAI2^V<u>F</u>LA^ also showed highly reduced responses to KAR_2_, but had stronger responses to KAR_1_ and *rac*-GR24 than wt AtKAI2. AtKAI2^VYIA^ had wt responses to KAR_1_, KAR_2_, and *rac*-GR24 (Figure 7B, S4B). AtKAI2<u>^LFLA^</u> seedlings showed the weakest rescue of hypocotyl elongation under mock-treated conditions, suggesting that KL perception may have been reduced more than in other variants (Figure S4B).

From these data, we conclude that Y124F is sufficient to reduce KAR_2_ response. Indeed, every transgene with Y124F conferred little or no response to KAR_2_. V96L also reduced KAR_2_ response as a single substitution, but did not have a consistent effect when combined with other substitutions. The effects of L139I and A161V substitutions were harder to decipher, but it is notable that five of the six variants with A161V had the weakest responses to *rac*-GR24.

## DISCUSSION

We identified two receptors, LsKAI2a and LsKAI2b, encoded in the lettuce genome that might activate germination in the presence of nanomolar KAR_1_. We propose that *LsKAI2b* is responsible for germination responses to KAR_1_ because 1) it is expressed more abundantly in achenes during early stages of imbibition than *LsKAI2a*, and 2) it confers sensitive and specific responses to KAR_1_ when expressed in *Arabidopsis thaliana*, while *LsKAI2a* does not. In the future, isolating loss-of-function mutations of these genes would be an ideal approach to conclusively determine their roles in KAR_1_-induced germination of lettuce. Interestingly, we found that a KAR_1_-responsive fire-follower, *Emmenanthe penduliflora*, also encodes two KAI2 proteins in its genome. EpKAI2a is similar to LsKAI2a in terms of the predicted volume and morphology of its ligand-binding pocket. By comparison, LsKAI2b and EpKAI2b are predicted to have substantially enlarged pockets and share similar amino acid substitutions at a small set of pocket positions. We hypothesize that EpKAI2b enables KAR_1_ perception during seed germination of *E. penduliflora*, but this will require further investigation.

We investigated how LsKAI2b mediates sensitive and specific responses to KAR_1_. By comparing the amino acid sequences of known (i.e. LsKAI2b and ShKAI2i) and putative (i.e. EpKAI2b) KAR_1_-responsive proteins to evolutionarily conserved paralogs from the same species that do not confer KAR_1_ responses (i.e. LsKAI2a, ShKAI2c, PaKAI2c, and putatively EpKAI2a), we surmised that the pocket residues at positions 96, 124, 139, and 161 may influence ligand-specificity. We performed a broader comparison of KAI2 sequences from 199 asterids, including 173 eu-asterid (lamiids and campanulids) species and 26 outgroup species from the Ericales and Cornales. We found that the vast majority of sequences fall into two groups that are distinguished by the residue at position 124. The 173 asterid species comprise 115 lamiids; of these, 105 (91.3%) have a KAI2 sequence with Y124, while 79 (68.7%) have a KAI2 with F124. Of 58 campanulid species, 57 (98.3%) have a KAI2 sequence belonging to the Y124 group, but only 7 (12.1%) have a KAI2 from the F124 group. Similarly, 100% of outgroup species are represented in the Y124 group, but only 2 outgroup species (8%) are represented in the F124 group. Amino acid identities at positions 96, 139, 161, and 190 were also well conserved within each group, but different across the groups. Thus, consensus combinations of V96, Y124, L139, A161, and A190, or L96, F124, I139, V161, and F190 are observed among asterid KAI2 proteins. The residues at these positions may have co-evolved (Figure 6). We do not yet know whether these combinations of amino acids evolved convergently among KAI2 paralogs in distinct lineages, or whether F124-type KAI2 paralogs were lost on many occasions within the asterids. However, the unequal representation of lamiid, campanulid, and outgroup species in the F124 group is suggestive of one KAI2 duplication shared among Lamiids, with independent, smaller-scale duplications in the campanulids and Ericales.

### Pocket residues that influence ligand specificity of KAI2

Many studies have used structural comparisons, molecular dynamics modeling, and *in vitro* biochemical assays to identify residues that may influence the ligand-specificities or affinities of KAI2 proteins, particularly with regard to SL perception (Nelson, 2021). Three recent studies, however, have examined how specific residues affect KAI2 ligand preferences *in vivo*. Unlike its relative in the Brassicaceae, *Arabidopsis thaliana*, the invasive, smoke-responsive species *Brassica tournefortii* is more sensitive to KAR_1_ than KAR_2_. Three *KAI2* genes are present in the *B. tournefortii* genome, only two of which appear to encode functional proteins (Sun et al., 2020). *BtKAI2b* is the most highly expressed *KAI2* paralog in seeds and seedlings. A *BtKAI2b* transgene confers stronger responses to KAR_1_ than KAR_2_ when expressed in Arabidopsis *kai2*, whereas *BtKAI2a* confers stronger or similar responses to KAR_2_ than KAR_1_. Swapping amino acid identities of BtKAI2a and BtKAI2b at positions 96 and 189 (per numbering in this study) switches their KAR preference *in vivo* and ability to respond to GR24^*ent*-5DS^ *in vitro*. Of the two residues, position 96 is primarily responsible for determining KAR_1_ versus KAR_2_ preference. BtKAI2c has an unusual R96 residue that makes the protein unstable (Sun et al., 2020). Its orthologs in other *Brassica* spp., which have a combination of L96, F124, L139, and V161 residues, might mediate KL-specific responses similar to AtKAI2<u>^LFLV^</u> (Figure S4). If so, perhaps the loss of BtKAI2c activity made *B. tournefortii* germination more dependent on external cues.

The legume *Lotus japonicus* also has two *KAI2* paralogs that show different ligand specificities. LjKAI2a responds similarly to KAR_1_ and KAR_2_, and responds better to GR24^*ent*-5DS^ than GR24^5DS^ (Carbonnel et al., 2020b). By contrast, LjKAI2b responds to KAR_1_, has very little response to KAR_2_, and does not respond to either GR24^5DS^ or GR24^*ent*-5DS^. Pocket residues at positions 157, 160, 190, and 218 differ between LjKAI2a and LjKAI2b. An unusual Trp substitution for Phe at position 157 is primarily responsible for the reduced responses to GR24^*ent*-5DS^ in LjKAI2b, although positions 160 and 190 may also contribute to a minor degree. The basis for different KAR responses has not yet been explored, or the role of position 218 (Carbonnel et al., 2020b). It is notable that LjKAI2a and LjKAI2b share V96, Y124, L139, A161 identities, and therefore would not have been anticipated to have different ligand specificities based upon our analysis.

A third study examined the basis of SL perception by KAI2d proteins from parasitic plants. A KAI2d paralog from *Striga hermonthica*, ShHTL7, confers exceptionally sensitive germination responses to SL when expressed in Arabidopsis (Toh et al., 2015). 92 variants of AtKAI2 were analyzed that had single, double, or triple substitutions for ShHTL7 residues at pocket positions 26, 124, 142, 153, 157, 174, 190, and 194 (Arellano-Saab et al., 2021). The variant proteins that showed the strongest yeast two-hybrid interactions with a MAX2 fragment in the presence of *rac*-GR24 tended to have substitutions at positions 124, 157, or 190. In transgenic Arabidopsis lines, however, the strongest germination response to *rac*-GR24 was conferred by a variant with W153L, F157T, and G190T substitutions. This variant gained responsiveness to 2’*R*-configured GR24^5DS^, while retaining responses to KAR_2_ and putatively KL (Arellano-Saab et al., 2021).

It seems likely that multiple combinations of pocket residues can produce similar ligand specificities in KAI2 proteins. For example, the KAR_1_-responsive proteins ShKAI2i and LsKAI2b differ from the consensus for F124-type KAI2 at position 139 (both are Val), and LsKAI2b also has M96 and A190 residues. We found that position 124 is an important determinant of KAR_2_ responsiveness, but others showed that position 96 influences this (Sun et al., 2020). This poses challenges for forming a predictive model of KAI2 ligand-specificity based upon amino acid sequences alone. It also implies that there are multiple evolutionary paths to produce convergent outcomes for KAI2 signaling. Several positions in KAI2 may be hotspots for the diversification of ligand preferences. In particular, residues at positions 96, 124, 157, 160/161, and 189/190 have been implicated in ligand selectivity by multiple studies (Sun et al., 2020; Carbonnel et al., 2020b; Arellano-Saab et al., 2021).

Potential benefits to agriculture may be achieved by understanding how to engineer KAI2 ligand preferences. KAI2 variants could be introduced to crops as transgenes, or endogenous KAI2 genes could be altered *in situ* through CRISPR-Cas9-based technologies. This could then allow selective activation of KAI2 signaling, which controls diverse traits, through application of a synthetic KAI2 agonist. We were not able to fully recreate the ligand selectivity of LsKAI2b in AtKAI2. Although we succeeded in removing KAR_2_ response while maintaining or enhancing KAR_1_ response, *rac*-GR24 response remained intact. One possible cause is that we performed substitutions at positions 96, 124, 139, and 161 with the consensus residues for F124-type KAI2 in asterids, which differ slightly from LsKAI2b identities.

### Evolution of smoke-induced germination

Several changes can be imagined to lead to the KAR_1_-dependent germination response observed in some fire followers. First, a KAR_1_ receptor may evolve more robust signaling activity. This could occur through an increase in affinity for KAR_1_ (or rather, a presumed KAR_1_ metabolite). Alternatively, enhanced affinity of the receptor for its signaling partners upon activation may increase signal transduction. This appears to be the case for ShHTL7 (Wang et al., 2021). Although *ShHTL7* transgenic lines respond to picomolar SL, the micromolar affinity of ShHTL7 for SL *in vitro* is comparable to other KAI2/HTL paralogs in *Striga hermonthica* (Toh et al., 2015; Tsuchiya et al., 2015; Wang et al., 2021). In contrast to other KAI2/HTL, ShHTL7 shows unusually high affinity for MAX2, which can be attributed to differences at five or fewer amino acids (Wang et al., 2021). It is unknown if ShHTL7 also has higher affinity for SMAX1 than other KAI2/HTL proteins. Second, increased expression of a KAR_1_-responsive receptor in seed may enable better germination responses to KAR_1_. As an example, we observed that *LsKAI2b* is more highly expressed than *LsKAI2a* in lettuce achenes (Figure 3). Similarly, the KAR_1_-preferring receptor *BtKAI2b* is more highly expressed in *B. tournefourtii* seed than other *KAI2* paralogs (Sun et al., 2020). Third, if KARs are metabolized *in vivo* as hypothesized, increased expression or activity of an enzyme(s) involved in that process could increase the availability of bioactive signals that activate KAI2. Fourth, an increase in physiological dormancy may be required to make seed germination more strictly dependent upon KAI2 signaling. One way this might occur is through downregulation of gibberellin biosynthesis or signaling. Arabidopsis germination typically requires gibberellins, which counteract the dormancy-promoting effects of abscisic acid. However, loss of SMAX1 through mutation or KAI2-SCF^MAX2^ activity can bypass this requirement (Bunsick et al., 2020).

## MATERIALS & METHODS

### Materials and plant propagation

KAR_1_, KAR_2_, and *rac*-GR24 were synthesized and provided by Dr. Gavin Flematti and Dr. Adrian Scaffidi (University of Western Australia). Oligonucleotide primer sequences are described in Table S4. *Lactuca sativa cv*. Grand Rapids achenes were sourced from a commercial supplier (185C, Stokes Seeds). The *Arabidopsis thaliana* double mutant line *d14 htl-3* (here referred to as *d14 kai2*) was kindly provided by Dr. Peter McCourt (University of Toronto) and is previously described (Toh et al., 2014). *Arabidopsis thaliana* and *Emmenanthe penduliflora* plants were propagated in Sungro Professional Growing Mix under white light (∼110 µmol m^−2^ s^-1^; MaxLite LED T8 16.5W 4000k light-emitting diode bulbs) with 16 h light/8 h dark photoperiod at ∼21-24°C. Soil was supplemented with Gnatrol WDG, Marathon (imidacloprid), and Osmocote 14–14–14 fertilizer.

### Functional analysis of *KAI2* genes from lettuce

The two *KAI2* sequences from lettuce were obtained from the Lettuce Genome Resource (Reyes-Chin-Wo et al., 2017) through a TBLASTN search of the *L. sativa* genome (version 4) using the *Arabidopsis thaliana* KAI2 (AtKAI2) protein sequence as a query. Each predicted protein sequence from lettuce was then used as a query in a reciprocal TBLASTN search of *Arabidopsis thaliana* (Araport 11) transcripts (Berardini et al., 2015) to identify likely *AtKAI2* orthologs. Lsat_1_v4_lg_4_361560640..361561729 was designated *LsKAI2a* and Lsat_1_v4_lg_4_361607540..361608739 was designated *LsKAI2b*. Each *KAI2* paralog was amplified from lettuce genomic DNA using primers with Gateway attB adapters, and Gateway cloning was used to shuttle each paralog via an entry vector into a plant binary destination vector (pKAI2pro-GW) that expresses genes under the control of the Arabidopsis *KAI2* promoter (Waters et al., 2015). Constructs were transformed into *Agrobacterium tumefaciens* (GV3101 pMP90), and floral dip transformation of *Arabidopsis thaliana* was performed in 5% sucrose (w/v) with 0.025% (v/v) Silwet-77 (Clough and Bent, 1998). Germination and hypocotyl assays were performed in a HiPoint DCI-700 LED Z4 growth chamber.

### Lettuce germination assays

Two layers of Whatman #1 filter paper (7 cm) were soaked with 2.5 mL of a treatment solution in a Petri dish. All aqueous solutions of KAR_1_, KAR_2_, and *rac*-GR24 were freshly prepared from 1000X stocks in acetone stored at −20°C. Approximately 50 lettuce achenes were plated onto each dish in the dark. Petri dishes were sealed with Parafilm and immediately placed into a growth chamber. Plates were incubated at 20°C for 60 min in dark, 10 min in far-red light (730 nm, 26 µmol m^-2^ s^-1^), and 47 h in dark. Germination was indicated by the emergence of a radicle.

### Gene expression analysis

Lettuce achenes were plated and light-treated as described for germination assays. Achenes were collected after 6 h or 24 h of imbibition and flash-frozen in liquid nitrogen before storage at −80°C. RNA extraction was performed with Spectrum Plant Total RNA Kit (Sigma). DNase I (New England Biolabs) digestion to remove contaminating genomic DNA was performed after RNA extraction. RNA concentrations were measured using a Qubit RNA Broad-Range Assay Kit (Invitrogen) and fluorometer. First-strand cDNA synthesis was performed with the Verso cDNA Synthesis Kit (Thermo-Fisher) with random hexamer and anchored oligo dT primers. Real-time quantitative PCR was performed on cDNA with Luna Universal qPCR Mastermix (New England Biolabs) in a CFX384 thermal cycler (Bio-Rad). Amplification conditions were 95°C for 3 minutes, and 40 cycles of 95°C for 15 seconds and 60°C for 1 minute, followed by a melt-curve analysis. *Arabidopsis thaliana* seedlings were grown as described for hypocotyl elongation assays and harvested at 5 days old. Gene expression analysis was performed as described for lettuce, except an On-Column DNase I Digestion kit (Sigma) was used during RNA extraction.

### Phylogenetic analysis of *KAI2*

The two *KAI2* sequences from lettuce were obtained as described above. Additional *KAI2* sequences from plant species representing the diversity of dicots, with *Physcomitrium* (*Physcomitrella*) *patens* as an outgroup, were collected from prior publications (Conn et al., 2015; Tsuchiya et al., 2015; Lopez-Obando et al., 2016; Yoshida et al., 2019). In total, 176 sequences from 56 species were combined, aligned, and manually adjusted with respect to predicted amino acid sequence. The nucleotide alignment was trimmed at the 5’ and 3’ ends to minimize gaps and regions of ambiguous alignment. Relative to the *AtKAI2* coding sequence, the final alignment retained codons 7 - 266 and consisted of 819 characters. The alignment was used to generate a Bayesian phylogeny in MrBayes v3.2.5 (Ronquist and Huelsenbeck 2003) (Ronquist and Huelsenbeck, 2003) as previously described (Conn et al., 2015)

### Hypocotyl Assays

Seeds were surface-sterilized (5 min in 70% EtOH with 0.05% (v/v) Triton X-100, followed by 70% and 95% EtOH washes, and air drying) and plated on 0.5x Murashige-Skoog (MS) Medium with MES Buffer and Vitamins (Research Products International), pH 5.7, solidified with 0.8% (w/v) Bacto agar (BD) supplemented with 0.1% (v/v) acetone or 1 µM KAR_1_, KAR_2_, or *rac*-GR24. Plates were stratified 3 d in dark at 4°C, then incubated in a growth chamber at 21°C for 3 h in white light (∼150 µmol m^-2^ s^-1^), 21 h darkness, and 4 d red light (660 nm, 30 µmol m^-2^ s^-1^). Seedlings were photographed and then hypocotyl lengths were measured using ImageJ (NIH).

### Structural Modeling and Analyses

Homology models of LsKAI2A, LsKAI2b, EpKAI2a, and EpKAI2b structures were generated with Phyre2 using ShKAI2iB (PDB structure 5DNW) as a template (Kelley et al., 2015; Xu et al., 2016). Structural illustrations were generated using PyMOL. Pocket volume and solvent accessible surface area were determined via CASTp (Dundas et al., 2006). Residues defining the pocket were broadly identified via CASTp and then probed for position and conservation to identify a final pocket-defining residue list to examine for all species.

### *Emmenanthe penduliflora* transcriptome Assembly

RNA was extracted from 7-d-old *Emmenanthe penduliflora* seedlings grown in 16:8 photoperiod on moistened filter paper using Spectrum Plant Total RNA kit (Sigma-Aldrich) after removal of seed coats. Library preparation was performed with 1000 ng of RNA input using NEB Ultra II Directional RNA kit with mRNA isolation. Sequencing was performed on a Nextseq 500 instrument with NextSeq mid-output 2×75 kit (paired-end 75 bp reads), producing 38.4 M reads (∼5.8 Gbp). The raw RNA-seq reads are available in NCBI Sequence Read Archive SRR16264938. A *de novo* transcriptome was assembled from paired-end reads with Trinity 2.6.6 (Grabherr et al., 2011) ran with the “--no_bowtie” parameter. Putative homologs of *KAI2* in *E. penduliflora* were identified by querying *AtKAI2* against the transcriptome assembly in a custom BLAST search and validated by reciprocal BLAST. *EpKAI2a* and *EpKAI2b* coding sequences are provided in Table S1.

### Analysis of KAI2 evolution in asterids

A reciprocal BLAST strategy was used to conservatively identify KAI2 orthologs. An AtKAI2 query was used in a BLASTP search of asterids in The 1,000 Plants Project Database (v5) (https://db.cngb.org/onekp/) (Carpenter et al., 2019; One Thousand Plant Transcriptomes Initiative, 2019). The 1000 best BLASTP hits were used in reciprocal BLASTP comparisons to Arabidopsis KAI2, D14, and DLK2. 164 BLAST matches with a match length less than 230 aa were filtered out as incompletely assembled genes or pseudogenes. 21 proteins that had potentially ambiguous orthology based on a difference of less than 50 in BLASTP bit scores between the first and second best hits to Arabidopsis proteins were also removed. Two KAI2 from a “*Mydocarpus* sp.”, presumably mislabeled, were removed. The remaining 813 asterid proteins were composed of 352 KAI2, 257 D14, and 204 DLK2. Multiple transcriptome assemblies were present for some species. 18 duplicates of KAI2 sequences from the same species were removed, leaving 334 KAI2 from 199 species. KAI2 were classified as Y124- or F124-type based upon the presence of an Ser-Pro-Arg-Tyr or Ser-Pro-Arg-Phe motif, which is highly conserved and bridges the intron splice junction. Only 12 proteins did not meet this criteria, and in some cases may be pseudogenes or incorrectly assembled. Asterid KAI2 protein sequences are provided in Table S2. Multiple sequence alignments of KAI2 proteins were performed in MEGA X and frequency plots of consensus sequences at pocket positions were visualized with WebLogo (Crooks et al., 2004; Kumar et al., 2018).

### AtKAI2 variant generation and analysis

A binary plant transformation plasmid expressing an *AtKAI2* coding sequence under control of an *AtKAI2* promoter (*pKAI2pro-AtKAI2*) was modified (Waters et al., 2015). A set of five oligonucleotides ranging from 41 to 56 nt in length (Table S4) were designed to span a central portion of the *AtKAI2* sequence encoding amino acids 96, 124, 139, and 161 when ligated end-to-end. Wildtype and mutant versions of the oligonucleotides were synthesized and phosphorylated with T4 polynucleotide kinase. Different combinations of wt and mutant phosphorylated oligonucleotides were annealed to four bridging oligonucleotides that were each complementary to the ends of two phosphorylated oligonucleotides. T4 DNA ligase was used to join the adjacent phosphorylated oligonucleotides into a continuous single strand of DNA. The new strand was amplified with Phusion high-fidelity DNA polymerase and inserted through Gibson assembly (New England Biolabs) into p*KAI2pro-AtKAI2* that had first been linearized by PCR to drop out the central region being swapped and digested with DpnI to remove un-linearized, methylated plasmid from the reaction. Sanger-sequence-verified *pKAI2pro-AtKAI2* variant plasmids were introduced into the *d14 kai2* mutant background via floral dip transformation. Transformants were selected at the seedling stage with hygromycin. Lines with single T-DNA insertion events were brought to homozygosity and characterized.

### Statistical analysis

Statistical analysis was performed in Prism 9 (GraphPad). Post-hoc statistical comparisons were performed after ANOVA or two-way ANOVA. Box plots show the median, 25th percentile, and 75th percentile. Tukey whiskers on box plots extend 1.5 times the interquartile range beyond the 25th/75th percentile up to the minimum/maximum value in the data set. Outlier data beyond Tukey whiskers are shown as individual points.

## ACKNOWLEDGMENTS

We thank Dr. Gavin Flematti and Dr. Adrian Scaffidi (University of Western of Australia) for providing KAR_1_, KAR_2_, and *rac*-GR24; Dr. Winslow Briggs (Carnegie Institution for Science) for providing *Emmenanthe penduliflora* seed; and Elise Landsbergen for extraction of RNA from *E. penduliflora*. RNA-seq library preparation and sequencing was performed by the Genomics Core at University of California, Riverside (UCR) and *de novo* transcriptome assembly was performed on the High Performance Computer Cluster at UCR. Funding was provided by the National Science Foundation (NSF-IOS 1856741 to DCN; NSF-CAREER 2047396 and NSF-EAGER 2028283 to NS; NSF-CAREER 2046256 to DK; and NSF Research Traineeship (NRT) Program Grant DGE-1922642 “Plants3D” to SM) and the United State Department of Agriculture (NIFA Award 2017-38422-27135 to SM; Hatch project CA-R-BPS-5209-H to DCN; and Hatch project CA-R-BPS-5154-H to DK).

## FIGURE LEGENDS

**Figure S1.**
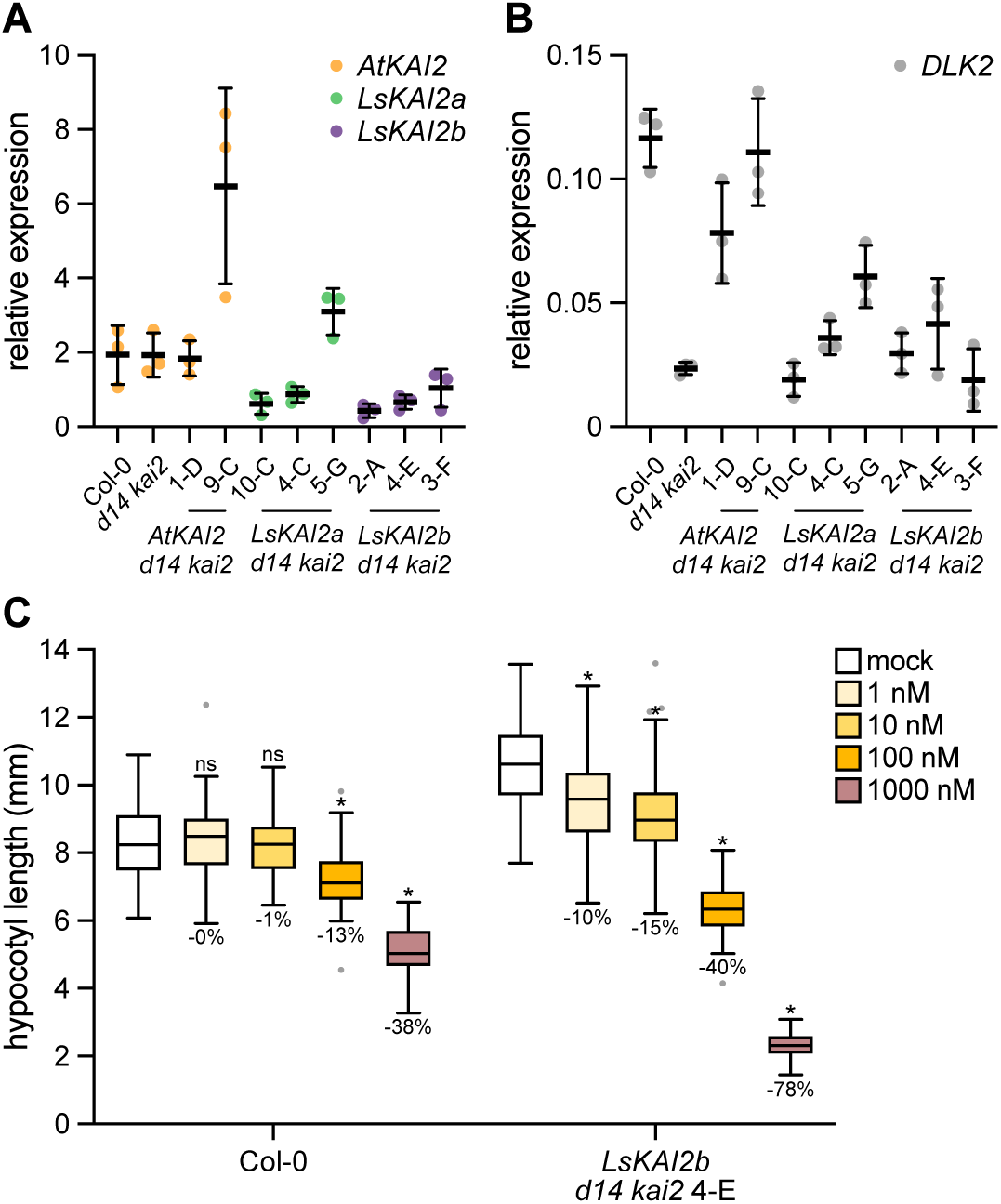
Transgene expression levels and KAR_1_ sensitivity in lettuce *KAI2* transgenic lines. A) qRT-PCR analysis of *AtKAI2, LsKAI2a*, and *LsKAI2b* expression, or B) *DLK2* expression relative to *CACS* in 5-d-old Arabidopsis seedlings grown under red light. n=3 pools of seedlings; mean ± SD. B) Hypocotyl length of 5-d-old Arabidopsis seedlings grown under red light in the presence of 1 nM to 1000 nM KAR_1_. Percent growth inhibition relative to mock-treated control within genotype is indicated below each treatment boxplot. *, p<0.01, Dunnett’s multiple comparisons test, treatment versus mock comparison within each line.

**Figure S2.**
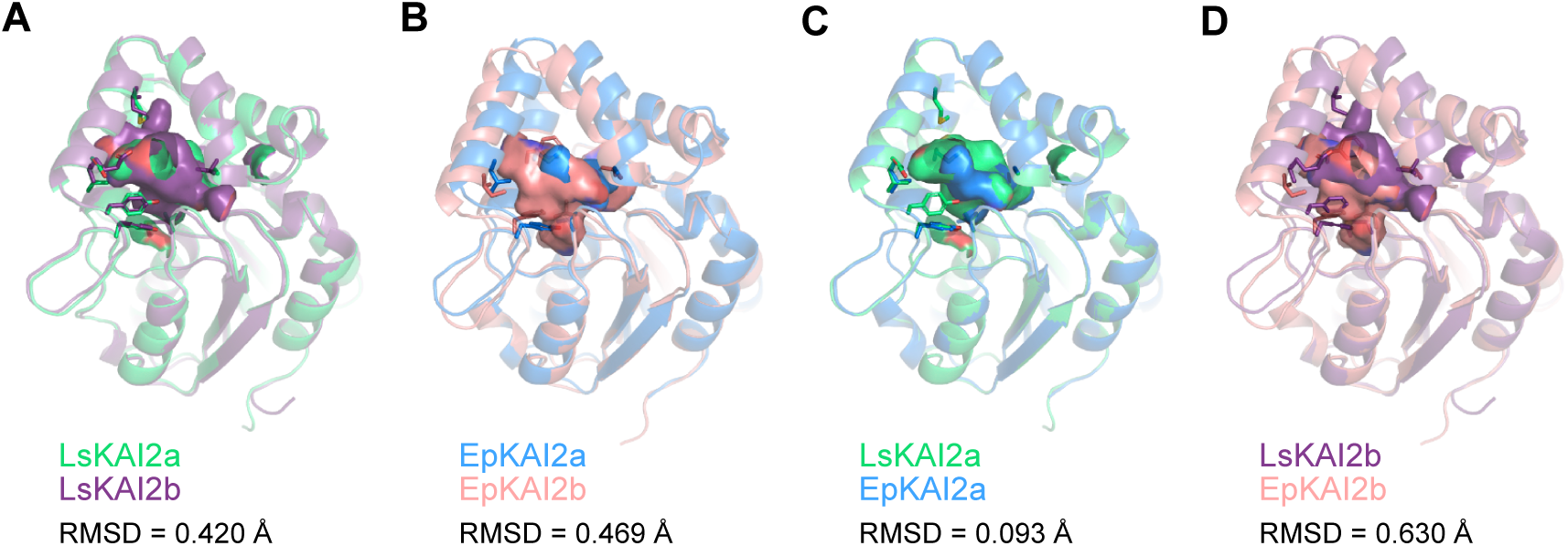
Comparisons of lettuce and *E. penduliflora* KAI2 structures within and across species. Overlaid homology models comparing A) LsKAI2a and LsKAI2b, B) EpKAI2a and EpKAI2b, C) LsKAI2a and EpKAI2a, and D) LsKAI2b and EpKAI2b. Hydrophobic cavities are shown with residues highlighted in Figure 6 as sticks. RMSD values are shown for each pair of models.

**Figure S3.**
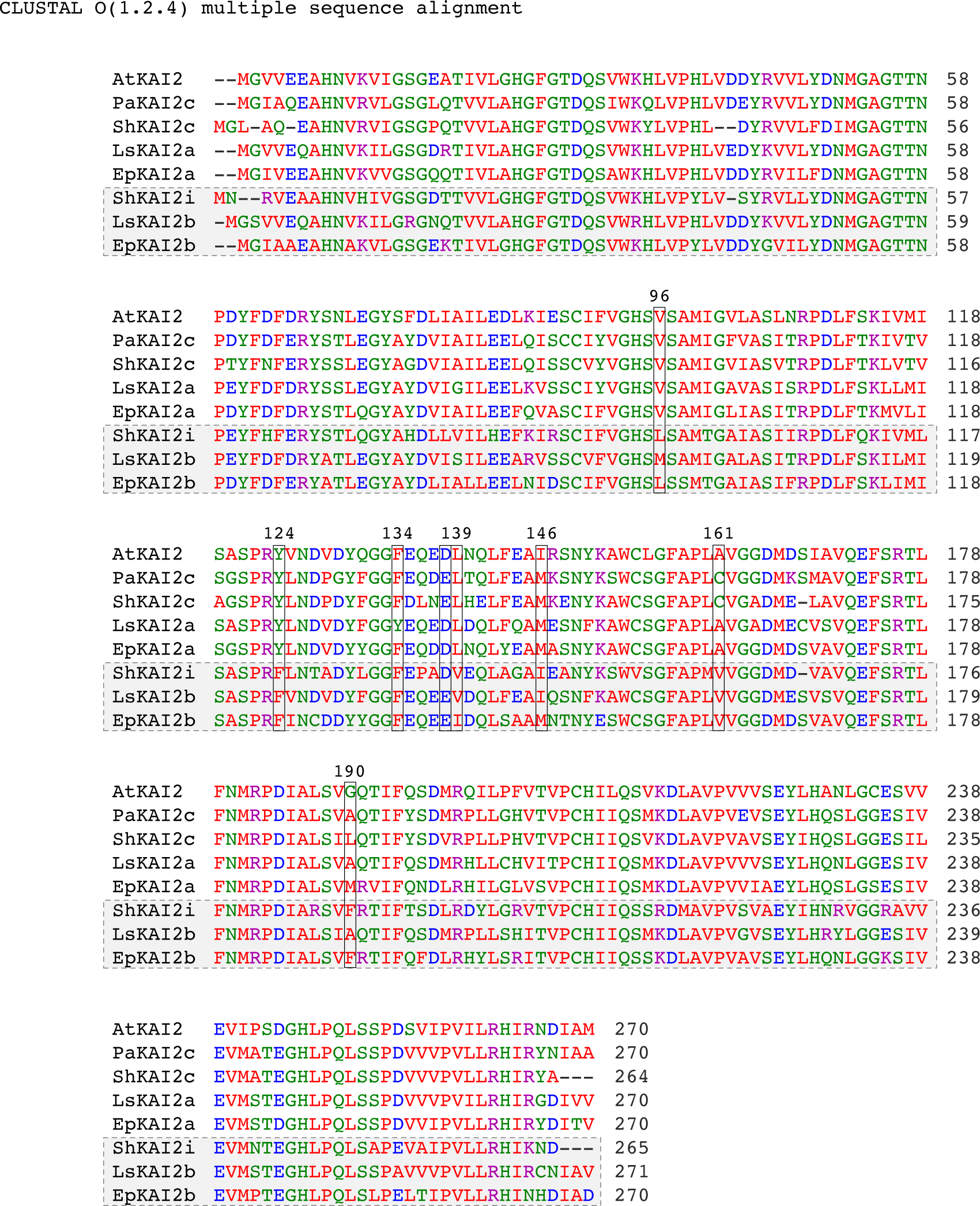
Sequence comparison of several characterized KAI2 proteins. Multiple sequence alignment by Clustal Omega of KAI2 proteins from *Arabidopsis thaliana, Phelipanche aegyptiaca, Striga hermonthica, Lactuca sativa*, and *Emmenanthe penduliflora*. ShKAI2i and LsKAI2b show selective responses to KAR_1_, and EpKAI2b is hypothesized to have similar properties. Pocket residues that were different between either LsKAI2a and LsKAI2b, or EpKAI2 and EpKAI2b are highlighted in boxes.

**Figure S4.**
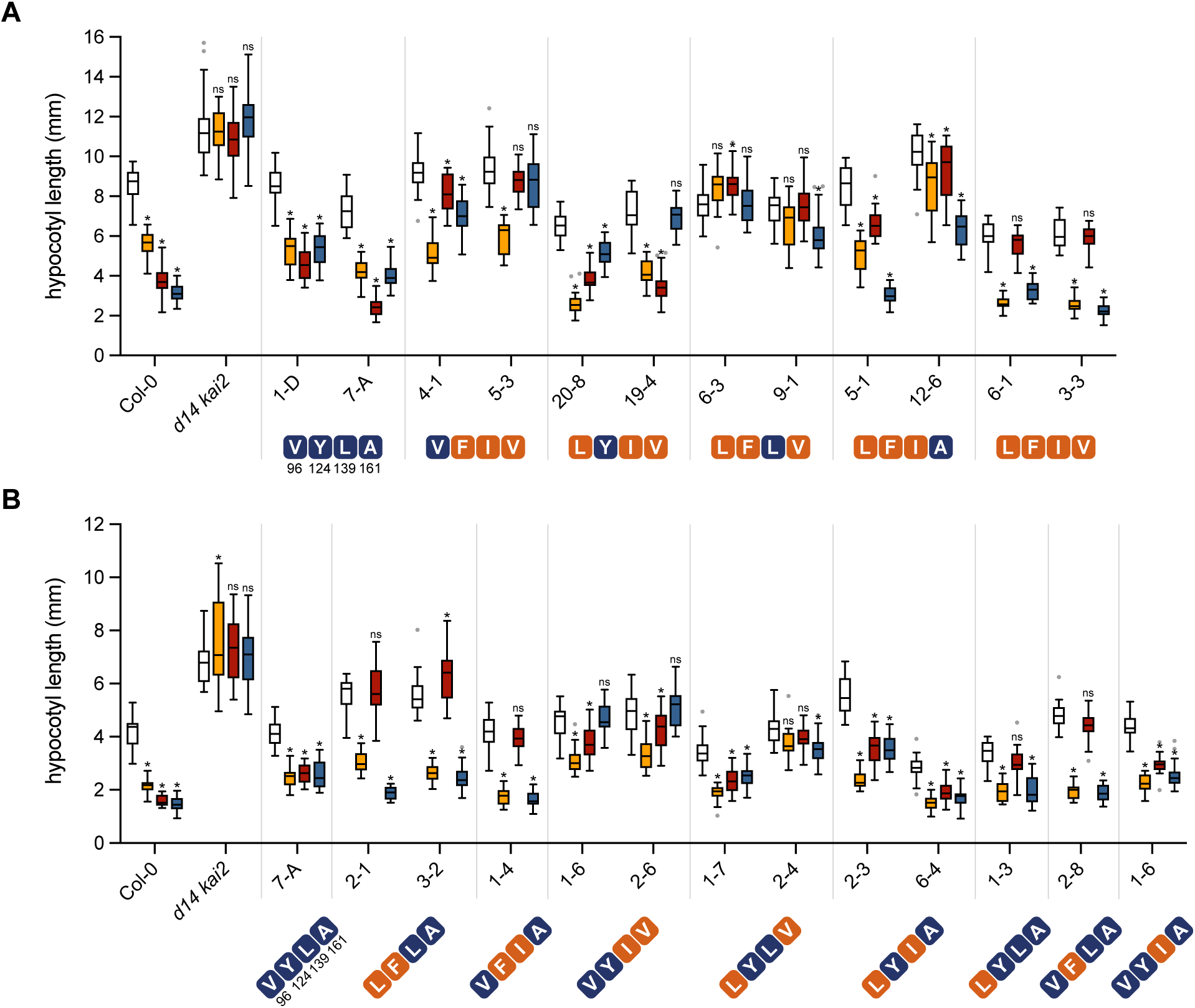
Hypocotyl elongation responses to KARs and *rac*-GR24 conferred by AtKAI2 variants. Hypocotyl lengths of 5-d-old Arabidopsis seedlings grown in red light in the presence of 0.1% (v/v) acetone or 1 µM KAR_1_, KAR_2_, or *rac*-GR24.Transgenic lines carry *AtKAI2* variants with A) quadruple- and triple-, or B) double- and single-substitutions at positions 96, 124, 139, and 161. All transgenic lines are in the *d14 kai2* double mutant background. n=20 (transgenics) or 40 seedlings (wt and *d14 kai2*) for A) and n=20 seedlings for B). Box plots indicate median and quartiles with Tukey’s whiskers. Gray dots indicate outlier data beyond Tukey’s whiskers. *, p<0.01, Dunnett’s multiple comparisons test, treatment versus mock comparison within each line. Growth inhibition responses to KAR and *rac*-GR24 treatments are summarized in Figure 7.

